# Mitochondrial Dysfunction Drives Age-Related Degeneration of the Thoracic Aorta

**DOI:** 10.1101/2025.06.13.659620

**Authors:** Arjune S. Dhanekula, Benjamin R. Harrison, Gavin Pharaoh, Aurora Mattson-Hughes, Stefano Tarantini, Rudolph Stuppard, Scott C. DeRoo, Christopher R. Burke, Billiana Hwang, Jay D. Pal, Michael S. Mulligan, David J. Marcinek

## Abstract

This study investigated the role of mitochondrial function in aortic aging. As the aorta ages, it becomes stiffer and less compliant, increasing the risk of aneurysmal disease, hypertension, and diastolic dysfunction. Given the role of mitochondrial dysfunction in non-age related aortopathies and as a hallmark of aging, we investigated its contribution to the aging aorta. Both male and female young (5-6 month) and aged (24-25 month) C57Bl/6J mice received mitochondrial-targeted peptide elamipretide (ELAM; SS-31) for 8 weeks. ELAM restored complex II-linked respiration in aged mice to values seen in young mice, while also improving relative phosphorylative flux. ELAM treatment also reduced inflammatory MMP9 expression and elastin breaks in aged mice. Bulk RNAseq analysis revealed that ELAM treatment significantly affected the aortic transcriptome in an age-dependent manner, reducing the expression of senescent and associated pro-inflammatory genes. Mitochondrial dysfunction thus drives aortic aging and is a potential therapeutic target for future study.

**Teaser:** Loss of mitochondrial function with age drives degeneration of the aging aorta, and thus is a potential target for therapy.

## Introduction

Aging is the greatest risk factor for cardiovascular disease, which continues to not only be the number one cause of mortality worldwide, but also a significant detriment to healthspan, quality of life, and public health spending[1]. Age-related changes in the thoracic aorta play a significant role in age-associated cardiovascular disease. Aortic compliance and structural integrity both decrease with age, not only increasing the prevalence of aortic pathology, such as aneurysm and dissection, but imparting significant physiological burden. These effects may directly contribute to diseases such as hypertension and heart failure with preserved ejection fraction[2–5]. The molecular mechanisms underlying aging in the aorta are not well understood, limiting the potential targets for clinically-approved therapies.

Mitochondrial dysfunction is a classic hallmark of aging[6]. Impairments in oxidative phosphorylation with age result in altered bioenergetics and redox homeostasis in multiple tissues[7–13]. The resulting rise in reactive oxygen species (ROS) and alteration of mitochondrial metabolism can trigger senescence through p16^INK4A^ and p53/p21^WAF1^ signaling[8]. Increased ROS production also contributes to genomic instability and inflammasome activation[8].

Mitochondrial dysfunction has been shown to contribute to chronic, age-related disease in many organs, including the brain, heart, kidney, and skeletal muscle[11, 12, 14–22]. Mitochondrial dysfunction is common in pathologic aortic tissue (including in patients and animal models with genetic aortopathies), but there is limited data demonstrating this in aged thoracic aortic tissue[7, 23–30].

Elamipretide (ELAM; SS-31) is a mitochondrial-specific peptide that has been shown to interact with cardiolipin in the mitochondrial membrane to restore the cristae membrane structure, improve the assembly and function of the electron transport system, and directly interact with mitochondrial proteins responsible for metabolism and signaling[11, 12, 14–20, 24, 31–34].

Administration of ELAM has resulted in treatment of age-related dysfunction in several organ systems, including a reversal of diastolic dysfunction in the heart, attenuation of age-related cardiac post-translational modifications, improvement of exercise tolerance in skeletal muscle, and reduction in abdominal aortic aneurysm formation[11, 12, 14–20, 24, 35]. Treatment with ELAM also reduces some of the senescent burden seen in these organs with age and disease[14, 17, 18]. Consequently, this drug is currently involved in multiple ongoing clinical trials[36]. The effects of ELAM have yet to be tested in thoracic aortic tissue.

We hypothesize that mitochondrial dysfunction is a major driver of aging in the thoracic aorta. In this study, we characterize mitochondrial function, structural changes, and global expression in the aging aorta, and assess our hypothesis by testing whether ELAM can alleviate age-related deterioration in aortic function.

## Results

### The Thoracic Aorta Becomes Diseased with Age

To assess the effects of age on the mouse aorta, we compared the thoracic aortas of aged (24-27 mo) C57BL6/JNIA mice to those of young (6-8 mo) C57BL6/JNIA mice. In vivo assessments of aortic diameter were made via echocardiography, measuring the diameter at the aortic root, ascending aorta, mid-aortic arch, and distal aortic arch (Figure 1A). Consistent with prior literature, the root and ascending aorta in old mice is significantly larger in diameter compared to young mice (Figure 1B)[2, 37]. We also observed a significantly increased frequency of elastin breaks and increased collagen content in the aorta of older mice (Figures 1C-1E). Elastin in the aorta absorbs the energy from systolic ejection and transfers it back into the blood during diastole, resulting in laminar flow[38]. However, loss of elastin integrity results in collagen deposition, increased thickness of the arterial wall (especially in the tunica media), and enhanced aortic stiffness[37, 39, 40]. There was no gross evidence of aneurysm, dissection, or rupture in any of the aortas, regardless of age.

**Figure 1:**
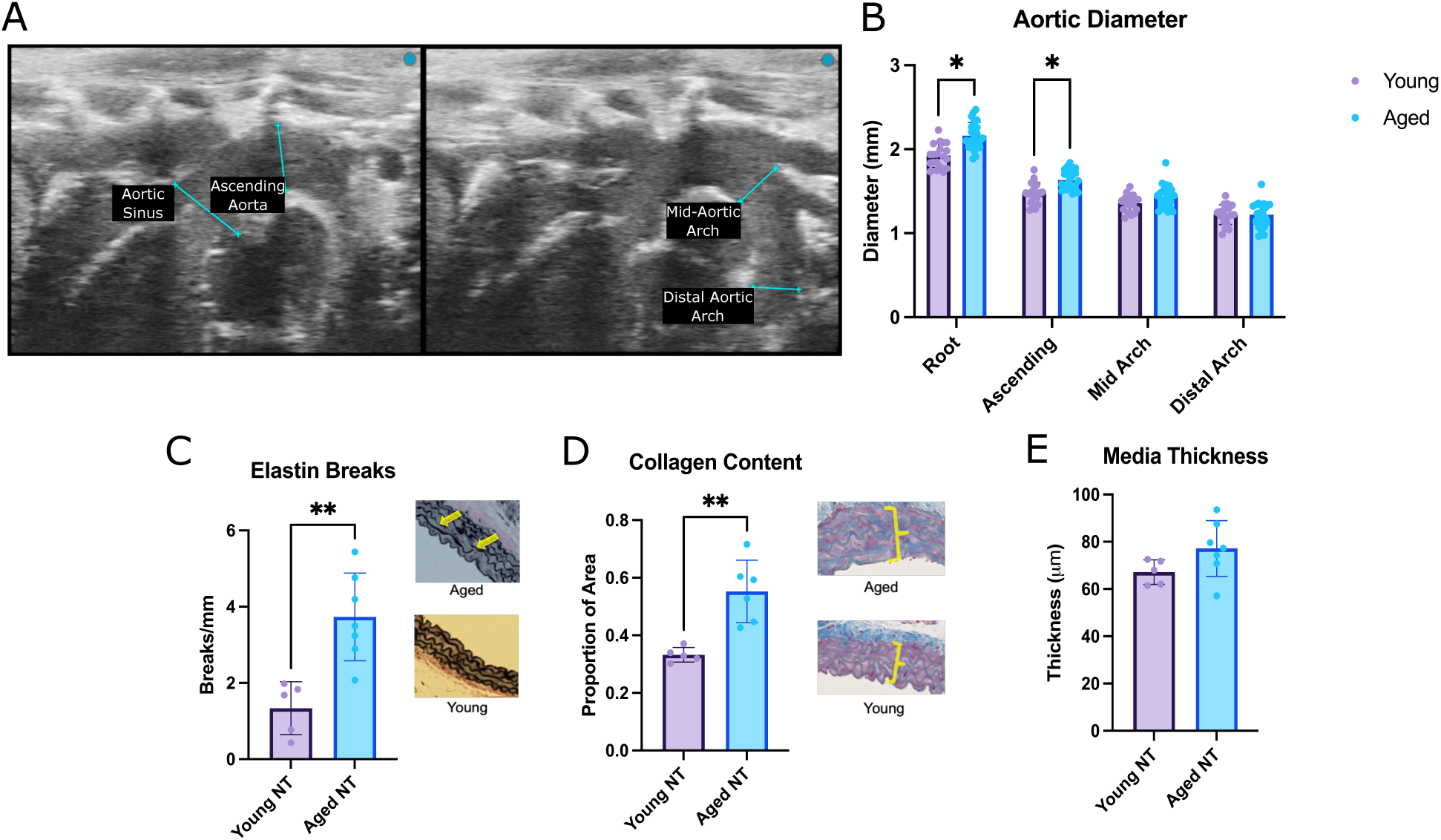
Age-related changes in the thoracic aorta. (A) Representative echo image of the murine aorta showing the aortic root, ascending aorta, mid-aortic arch, and distal aortic arch. (B) Comparison of maximal aortic diameter between young (6-8 mo, n=19) and aged mice (24-27 mo, n=25). Histologic comparisons of elastin breaks (C), collagen content (D), and media thickness (E) between young (n=5) and aged (n=6-7) mice. Multiple t-tests (with Holm-Sidak correction) and student’s t-test, * p<0.05, ** p<0.01, *** p<0.001, **** p<0.0001.

### Elamipretide Improves Complex II-Linked Respiration in the Aorta

We hypothesize that mitochondrial dysfunction drives the pathologic changes seen in the aging aorta. We thus assessed age-related changes in aortic respiration and made parallel measurements on aortas from mice treated for eight weeks with ELAM. In the ascending aorta and aortic arch, there is a significant decline in Complex II-linked respiration with age (Figures 2A-2E). However, treatment with ELAM ameliorated this drop in Complex II-linked respiration, with an associated significant increase in maximal coupled respiration (Figures 2A-2E). ELAM had no significant effect on respiration in the young ascending aorta/arch.

**Figure 2:**
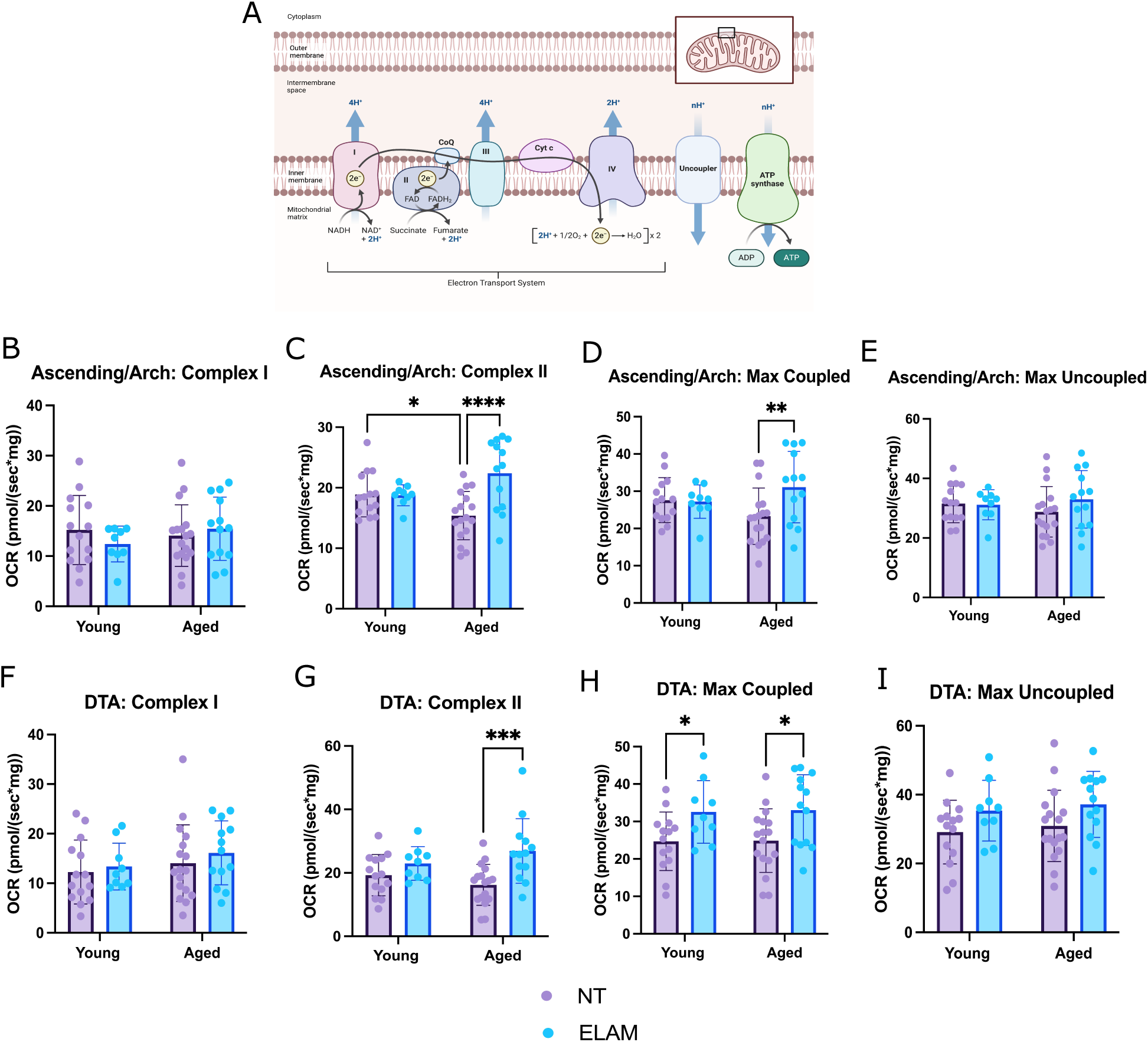
Complex II-linked respiration declines with age but is rescued with ELAM. (A) Schematic of the electron transport system; note the uncoupler, which breaks the link between aerobic respiration and ATP production. Oxygen consumption rate (OCR) of complex I (B), complex II (C), maximal coupled respiration (D), and maximal uncoupled respiration (E) in the ascending aorta and aortic arch of young NT (n=14), young ELAM (n=9), aged NT (n=17), and aged ELAM (n=13) mice. Oxygen consumption rate (OCR) of complex I (F), complex II (G), maximal coupled respiration (H), and maximal uncoupled respiration (I) in the descending thoracic aorta (DTA) of the same mice. 2-way ANOVA, Fisher’s least significant difference, * p<0.05, ** p<0.01, *** p<0.001, **** p<0.0001. (A) was created with BioRender.com.

In the descending thoracic aorta (DTA), there are no statistically significant differences in respiration between the aged and young DTA of untreated mice (NT) (Figures 2F-2I). However, treatment with ELAM significantly increased Complex II-linked respiration and maximal coupled respiration in the aged aorta. Treatment with ELAM also increased maximal coupled respiration in the young DTA, as well (Figures 2F-2I).

### Elamipretide Improves Phosphorylative Capacity in the Aged Aorta

Phosphorylative capacity (also referred to as the phosphorylation system control ratio) refers to the extent to which mitochondria can phosphorylate ADP to ATP at maximal coupled respiration [41, 42]. In young mouse aortas, phosphorylative capacity is nearly identical to the respiratory capacity of the electron transport system (maximal uncoupled respiration) (Figure 3). This suggests that the electron transport system is not limited by the rate of ADP phosphorylation.

**Figure 3:**
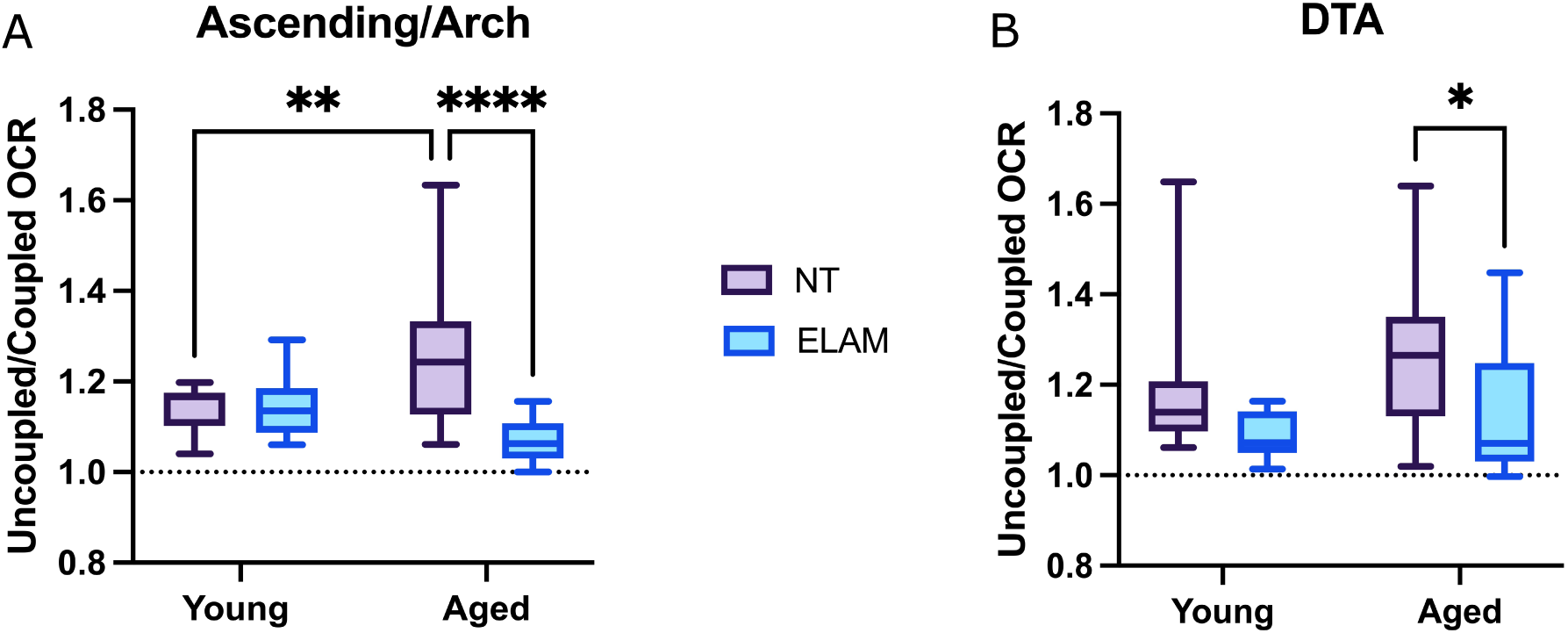
Phosphorylative capacity is rate-limiting with age and improves with ELAM treatment. Relative phosphorylative capacity in the ascending aorta and arch (A) and DTA (B). Young NT (n=14), young ELAM (n=9), aged NT (n=17), and aged ELAM (n=13). 2-way ANOVA, Fisher’s least significant difference, * p<0.05, ** p<0.01, *** p<0.001, **** p<0.0001.

However, in aged mouse aortas, especially in the ascending/arch, there is an increase in the ratio of uncoupled to coupled respiration, suggesting a deficiency in phosphorylative capacity that limits flux through the electron transport system (Figure 3A). Treatment with ELAM improved this deficit, reducing the uncoupled/coupled ratio closer to that in the aorta of young mice in both the aged ascending/arch (Figure 3A) and aged DTA (Figure 3B). There was no difference after ELAM treatment in the young aorta (Figures 3A-3B).

### Mitochondrial Dysfunction Induces MMP9 Expression with Age, but not Canonical Markers of Senescence

We have demonstrated anatomic and histologic changes in the aging aorta, along with a reduction in mitochondrial bioenergetic function. Treatment with ELAM, a mitochondrial targeted peptide, improved these bioenergetic changes. We then sought to understand the extent to which mitochondrial function might drive other changes in the aorta with age.

Cellular senescence is a state of replicative exhaustion, and increased senescent burden is a hallmark of aging. Senescent cells produce inflammatory signaling (the senescence-associated secretory phenotype, SASP), resulting in organ dysfunction[23, 43–53]. Furthermore, mitochondrial function is often a major driver of the senescent phenotype[8]. Thus, we hypothesize that treatment with ELAM would improve the senescent and inflammatory phenotype of the aged aorta.

To assess the level of senescence in the aged aorta, we measured the expression of the canonical senescence markers p16/CDKN2a (p16) and p21/CDKN1a (p21)[54]. The ascending/arch and DTA were pooled together for this analysis due to the small size of the murine aorta. Consistent with an increased senescent burden with age, transcript abundance of p16 and p21 were higher in the aged aortas relative to young aortas. ELAM treatment did not affect either p16 or p21 expression in aged mice, but did decrease p21 expression in young mice (Figure 4A-B).

**Figure 4:**
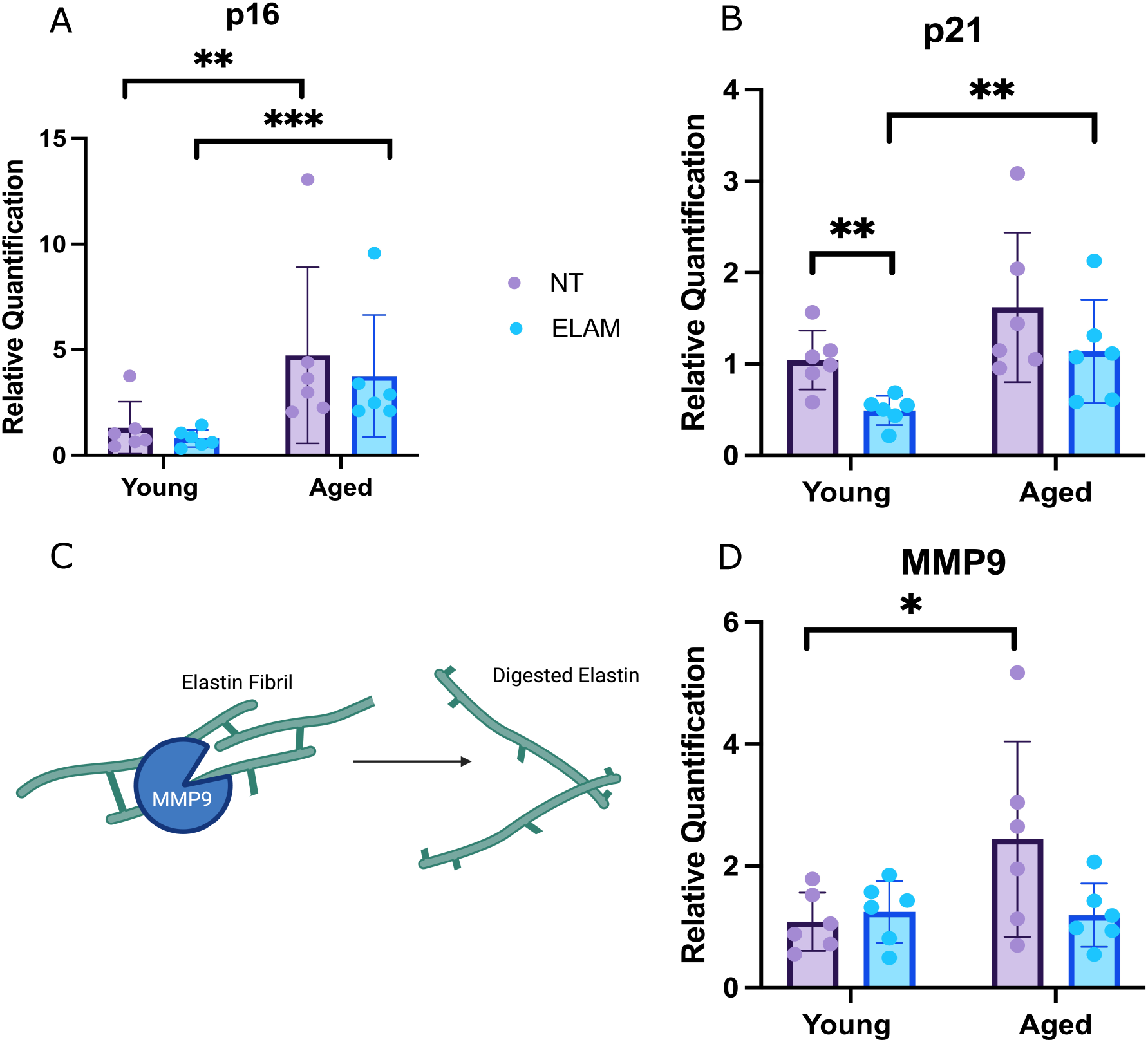
Canonical senescent markers and the inflammatory enzyme MMP9 increase with age, and ELAM treatment reduces MMP9 expression. RT-qPCR of the murine thoracic aorta for the senescent markers p16 (A) and p21 (B). (C) Schematic of MMP9’s mechanism of action in the vascular extracellular matrix. (D) Expression level of MMP9 in the murine thoracic aorta. n=6 for all groups. 2-way ANOVA, Fisher’s least significant difference, * p<0.05, ** p<0.01, *** p<0.001, **** p<0.0001. (C) was created with BioRender.com.

We found that that the expression of matrix metalloprotease 9 (MMP9)—a common SASP and inflammatory enzyme central to aortic wall degeneration—was significantly higher in aged NT mice when compared to younger mice (Figure 4C-D) [55–60]. However, the aged mice treated with ELAM showed no difference in MMP9 transcript levels relative to the young mice (Figure 4D).

### Mitochondrial Dysfunction Causes Elastin Fragmentation in the Aged Aorta

We then investigated whether mitochondrial function is important in preventing age-related degeneration of the aortic wall. We measured the aorta by echocardiography at the level of the root, ascending aorta, mid-arch, and distal arch both prior to treatment (pre) and at the end of the 8-week treatment period (post). We largely saw no effect of ELAM or age on aortic diameter over the study period (Figure 5A-B, Supplemental Figure 1A-B). The only difference was between aged NT (<1% decrease in diameter over the study period) and aged ELAM (5% increase in diameter over the study period) in the ascending aorta (Figure 5B). Evaluation of the left ventricle revealed no effect of ELAM on left ventricular ejection fraction, wall thickness, or chamber diameter (Supplemental Figure 1C-J). Blood pressure did not change over the treatment period for any of the groups (Supplemental Figure 2A-L).

**Figure 5:**
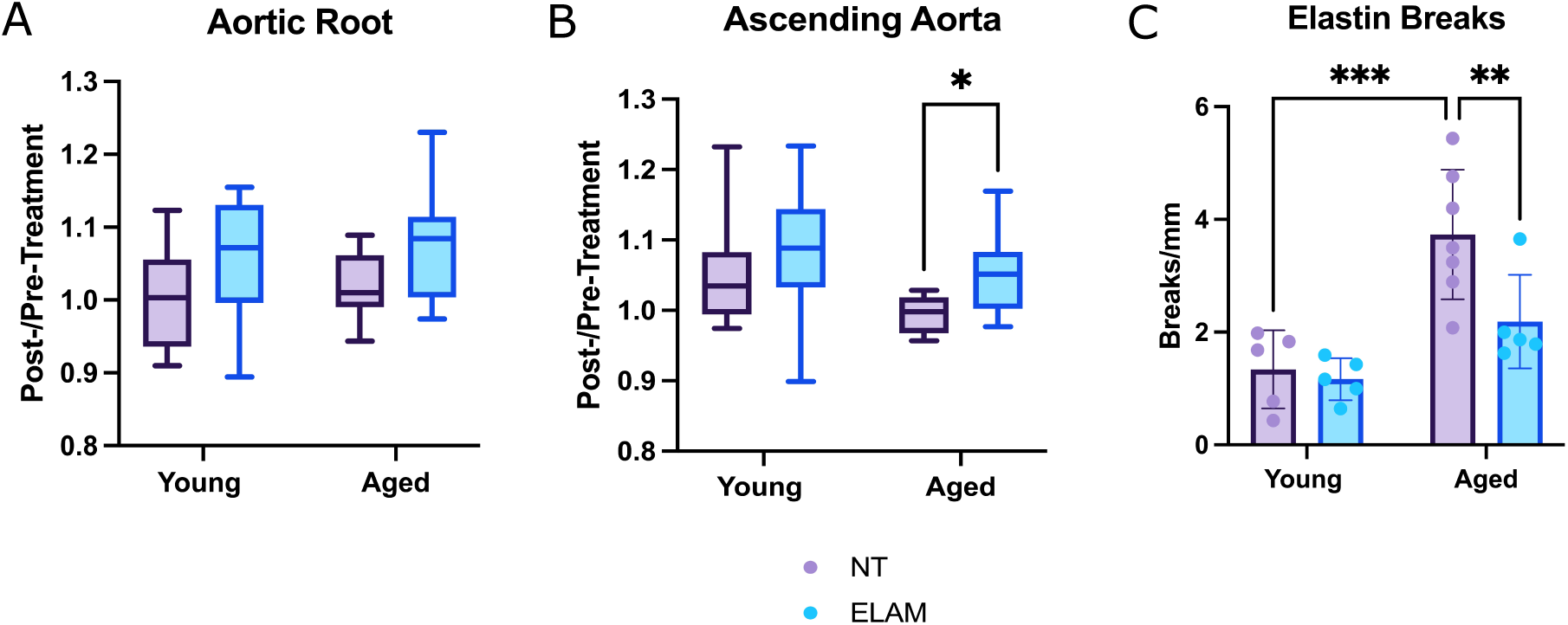
ELAM improves elastin breaks with age. Relative change of the aortic root (A) and ascending aorta (B) over the 8 week study period in young NT (n=10), young ELAM (n=9), aged NT (n=12), and aged ELAM mice (n=13). Histologic assessment of the frequency of elastin breaks (C) in the four treatments groups (n=5-6 per group) reveals a significant improvement in elastin breaks with ELAM treatment. 2-way ANOVA, Fisher’s least significant difference, * p<0.05, ** p<0.01, *** p<0.001, **** p<0.0001.

Histologically, the aged ELAM group did not differ in media thickness or collagen content relative to the aged NT group (Supplemental Figure 3A-B). However, treatment with ELAM did result in significantly fewer elastin breaks in the aged ELAM group relative to the aged NT group, and the older ELAM-treated mice had an elastin break frequency similar to young mice (Figure 5C). None of the harvested aortas (either ELAM or NT) had any grossly obvious aneurysms, dissections, or ruptures.

### Mitochondrial Dysfunction Drives Senescence and Inflammation in the Aging Aorta

The improvement in aortic histology seen in older mice treated with ELAM suggests that mitochondrial dysfunction may be a driver of the aged phenotype in the mouse aorta. To further investigate the extent to which mitochondrial dysfunction affects age-related changes in the aorta, we examined the transcriptome of the aortas of old and young mice following ELAM treatment compared to NT using RNAseq.

Among 24 samples we measured expression of 20,607 genes. Approximately 59% of the variance in the aorta transcriptome was captured by three principal components (PCs), with the first component (PC1) explaining 27% of variance. The transcriptome of aged NT mice differed substantially from the younger mice along PC1, whereas the transcriptome of NT mice differed to a lesser extent, if at all, across PCs 2 and 3 (Figure 6A). These data indicate that PC1 largely captures the effects of age in the mouse aorta. In stark contrast to aged NT mice, the transcriptome of older ELAM-treated mice along PC1 was similar to that of the younger mice (Figure 6A). To identify individual genes whose expression varies by age, ELAM, or an age-ELAM interaction, we then fit a linear model on the data. Investigating the effect of treatment revealed 6,287 differentially expressed genes, effect of age revealed 6,061 genes, and effect of the age-treatment interaction revealed 4,339 genes (Supplemental Table 1).

**Figure 6:**
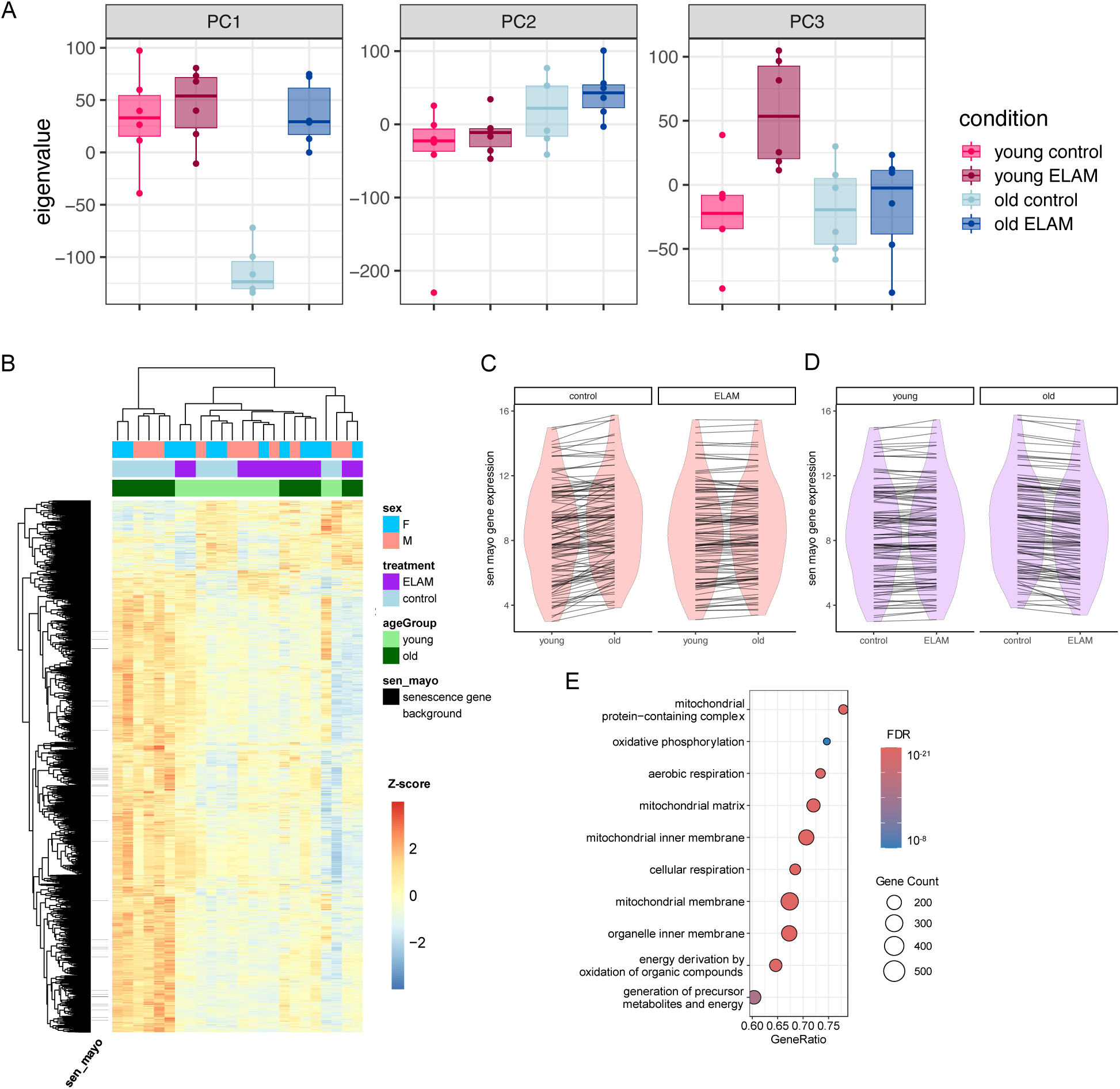
ELAM affects the aortic transcriptome in an age-dependent manner. (A) Principal component (PC) analysis of the aortic transcriptome across the four treatment groups (n=6 per group). (B) Heat map showing the clustered expression level of genes with significant age by treatment interaction (ANOVA, FDR<0.05). Note the SenMayo genes demarcated by black hashes on the left side, and the significant difference between the aged control (NT) mice and the other treatment groups. The mean expression level of each of the SenMayo genes in young and aged mice are connected by lines in (C), and the mean expression level of each of the SenMayo genes in ELAM and control mice are connected by lines in (D). (E) The top ten most significant GO terms enriched by genes with age-treatment interaction.

Genes with age-treatment interactions (i.e. the relationship between mitochondrial function and aging) are anticipated by the separation of aged NT mice from the aged ELAM mice along PC1. We anticipate that many of the genes with such effects would show a large difference in aged mice compared to the young NT mice, and that these effects of age would be tempered by ELAM treatment. Indeed, clustering the genes and samples by the expression of the 4,339 genes with an age-treatment interaction led to clear distinction of the aged NT mice compared to both the young mice and the ELAM-treated older mice (Figure 6B). The expression of a majority of these genes were upregulated in aged NT relative to young NT mice (Figure 6B, Supplemental Figure 4A). Treatment with ELAM led to a substantial difference in the expression of these genes in aged mice, with the majority of genes reverting toward expression that was similar to young mice (Figure 6B, Supplemental Figure 4B). ELAM had minimal effect on gene expression levels in young mice (Figure 6B, Supplemental Figure 4B). Thus, ELAM treatment has a more substantial effect on gene expression in aged aortic tissue than in young tissue.

We then utilized the SenMayo dataset to assess the effect of age and ELAM on senescence, as SenMayo is a validated gene set that can identify a broad range of senescent and associated inflammatory/SASP expression across multiple tissue types[61]. Genes in the SenMayo dataset are significantly enriched in both the 6,061 age-dependent genes and the 4,339 age-treatment interaction genes, but not in the genes with affected by treatment alone (Table 1). Thus, age affects senescent expression in the aorta and treatment with ELAM affects the expression of SASP in the aortas of older mice. Consistent with elevated senescence in the aged aorta, we see remarkably higher expression of SenMayo genes in the aorta of older mice (Figure 6C).

**Table 1:** SenMayo genes are enriched among the age- and age-dependent treatment genes. 56 and 43 of the 114 mouse genes in the SenMayo gene set are found among the 6,061 or 4,339 genes with treatment effects or age-dependent treatment effects, respectively[61]. Only 34 and 24 genes would be expected to intersect both model effects and SenMayo genes by chance, a significant enrichment. Fisher’s test, odds ratio >2.28, P < 1.1×10^−5^.

| Term | # genes affected (FDR <5%) | # SenMayo genes (n=114) | # expected by chance | p-value |
| --- | --- | --- | --- | --- |
| treatment | 6,287 | 31 | 35 | 0.477 |
| age (young vs. old) | 6,061 | 56** | 34 | $1.09 \times 10^{-5}$ |
| treatment x age | 4,339 | 43** | 24 | $4.31 \times 10^{-5}$ |

Furthermore, the aortas of older mice treated with ELAM have significantly lower SenMayo expression than aged NT mice (Figure 6D). These data suggest that mitochondrial dysfunction drives senescence and associated inflammatory expression in the aged aorta.

Gene ontology (GO) analysis of the genes with age-treatment interaction revealed 629 significantly terms (FDR<5%). The top ten GO terms by adjusted p-value were nearly all related to mitochondrial structure and function. The majority were related to electron transport chain and membrane proteins, except for two metabolic GO terms (organic acid metabolic process and oxoacid metabolic process) (Figure 6E). Amongst all the significant GO terms, mitochondrial function/assembly/biogenesis, metabolic pathways, and inflammatory pathways were frequently recurring terms (Supplemental Data 1).

## Discussion

Age-related changes in the aorta play a significant role in cardiovascular morbidity and offer a promising area for clinical translation. However, the mechanisms driving these changes are still not fully understood. In this study, we investigated the role of mitochondrial function in aortic aging using the targeted peptide elamipretide (ELAM; SS-31). We had three major findings: 1) as the aorta ages, it develops a reduced capacity for complex-II linked respiration and decreased relative phosphorylative capacity, but both are improved with ELAM treatment; 2) ELAM administration decreases the extent of age-related inflammatory destruction in the aortic wall; 3) aging increases the inflammatory and senescent transcriptome in the aorta, and ELAM reverses those changes. No sex differences were noted throughout the study. These results not only suggest that mitochondrial dysfunction is a driver of aging in the aorta, but that interventions even later in life could ameliorate aging physiology. Beneficial effects of late-age treatment have been seen with rapamycin and ELAM[35, 62]. Here we show that late-age ELAM treatment— and perhaps other interventions targeting the mitochondria—may serve as an important treatment for age-related vascular disease.

Alterations in mitochondrial bioenergetics are central to both aging and age-related disease. We identified a decrease in complex II-linked respiration and phosphorylative capacity in the aging aorta. Complex II has an important role in cellular bioenergetics, and mutations in this complex can produce hereditary neurodegenerative disorders like Leigh’s syndrome[63]. However, more relevant to this discussion is the role of complex II in age-related disease. Studies in rodents and human cell lines have shown that complex II has reduced function in the aging brain, heart, liver, and integumentary system; complex II dysfunction has also been described in neurologic disorders like Alzheimer’s[63]. Diminished ADP phosphorylation is also seen with age-related organ decline. Even in the setting of saturating substrates, efficient phosphorylation of ADP requires proper functioning of the electron transport system along with effective delivery of ADP through the adenine nucleotide translocator (ANT) into the matrix and to complex V (ATP synthase). In other organs, aging results in reduced efficiency of ADP phosphorylation, while treatment with ELAM improves this phosphorylative efficiency[12, 64–67]. Our study revealed similar age-related changes in both complex II and ADP phosphorylation in the thoracic aorta, with ELAM effectively improving these phenotypes.

Both the transcriptomic and histologic data obtained in this study reveal a significant increase in inflammatory-driven destruction with age (“inflammaging”), a phenomenon already well-described in other organ systems[59, 68]. In the aorta, this destruction manifests as elastin degradation (mediated through MMPs), collagen deposition, and aortic dilation[37, 39, 40, 55, 56]. In the abdominal aorta, the inflammation is tied to atherosclerosis[69]. In the thoracic aorta, however, inflammatory destruction is rarely associated with atherosclerosis; the mechanisms driving thoracic aortic inflammation are thus less clear[70, 71]. Our data reveal age-related changes in the mitochondria as a driver of age-driven inflammation in the thoracic aorta, at least partially mediated through an increased senescent burden[53, 72]. While the canonical senescent markers p16 and p21 were not altered with ELAM treatment, there is a growing body of literature showing that these markers can be limited in accurately assessing the senescent phenotype[61, 73–75]. Using a much more diverse gene set developed to address these problems, a consistent reduction in age-associated senescent expression (including the inflammatory SASP) is seen with ELAM treatment[61]. Thus, improvement of age-related changes in mitochondria, even late in life, reduced senescent expression and resultant inflammatory signaling, resulting in a more youthful histologic architecture of the aorta.

We thus propose a mechanism in which mitochondrial changes with age (indicated by bioenergetic derangements) contribute to increased senescence and inflammation, elastin degradation, and subsequent aortic wall degeneration (Figure 7). ELAM administration improved bioenergetic function in an age-dependent manner. This treatment caused the expression profile in the aged aorta to take on a more “youthful” appearance, reduce senescent expression, and improve inflammation. Concurrently, the ELAM-treated aged aortas had a significantly reduced burden of elastin breaks.

**Figure 7:**
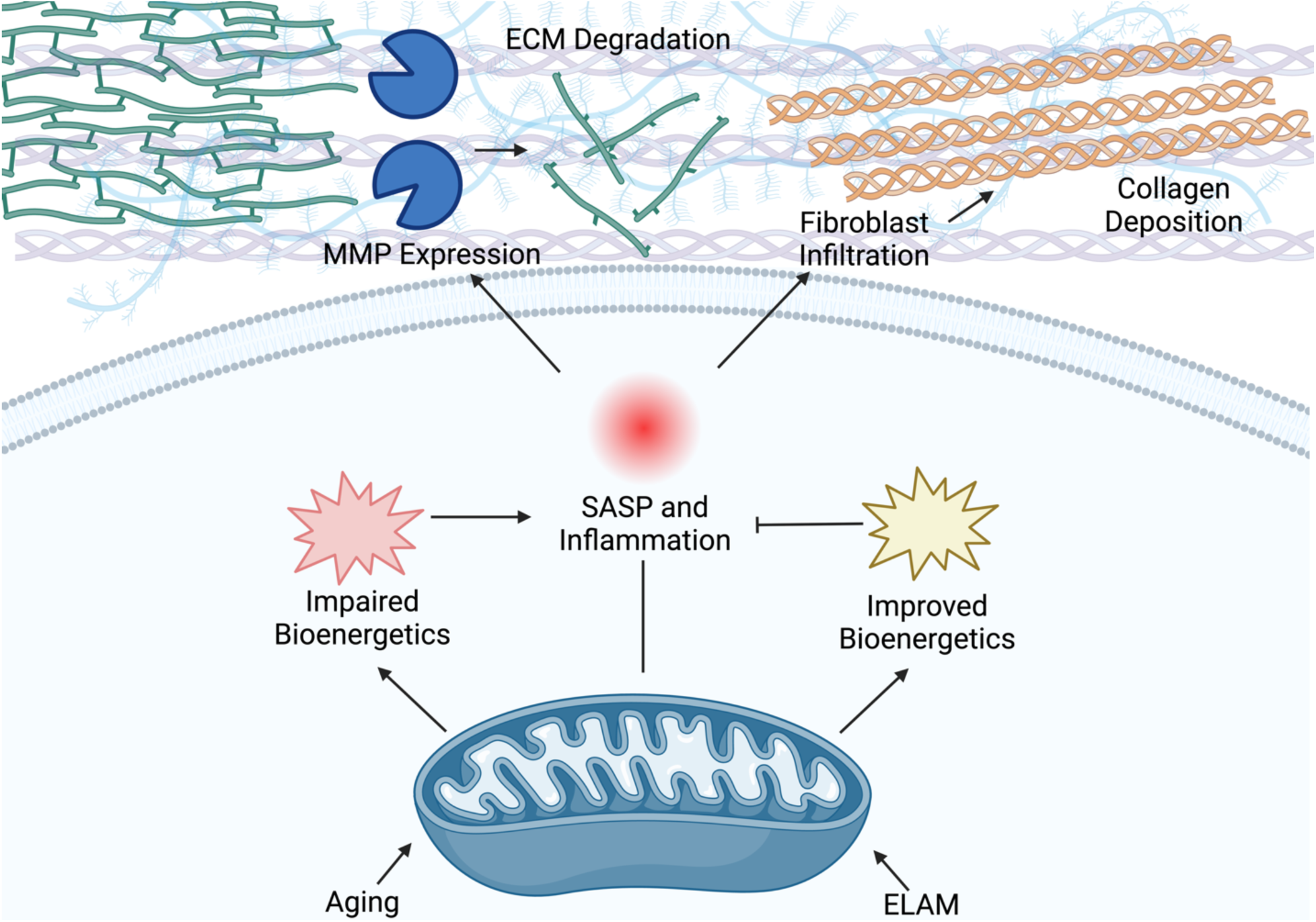
Proposed mechanism of age-related disease in the aorta. Bioenergetic derangements with age drive senescent expression and consequential inflammatory destruction of the aorta. ELAM blunts these inflammatory changes by improving the bioenergetic profile of the aging aorta. Created with BioRender.com.

This study had a number of limitations. First, while the mitochondrial-specificity of ELAM has been studied in detail, to our knowledge, this has not been confirmed in the aorta[31–34]. Thus, we cannot be certain that the effects of ELAM are due to direct effects on the mitochondria, although the top GO categories altered by the age-treatment interaction all point to mitochondrial related genes. Furthermore, the degree to which the expression-level and histologic changes seen after ELAM administration are dependent on the concomitant bioenergetic alterations of ELAM is also unclear. Future study would benefit from long-term administration and confirmation of the mitochondrial-specificity of ELAM in the aorta. Dynamic assessments of aortic contractility and compliance would also provide a more direct link for the effect of age and ELAM on physiologic changes in the aorta. Finally, proteomic data may provide a better surrogate for tissue function than transcription data.

Our data also revealed a relative increase of ∼5% in ascending aortic diameter over the treatment period in aged ELAM mice, while untreated aged mice had essentially no change in ascending aortic diameter. The other measured portions of the aorta also had no change in diameter amongst all treatment groups. A growth rate of 5mm/year in humans (∼10% relative growth) is an indication of aneurysmal disease[76]. Aneurysmal growth is often secondary to loss of elastin and increased stiffening of the aorta, reducing the ability to absorb systolic forces from the ejecting heart[37–40, 77]. However, the reason for the change in ascending aorta diameter in the ELAM treated aged mice is unclear; given the improvement in aortic wall architecture and elastin integrity with ELAM treatment, this increase in size is likely not due to inflammatory destruction of the wall. This apparent paradox warrants further investigation, as ELAM may be affecting biomechanical properties of the aorta in ways this study didn’t address. Future experiments directly investigating aortic compliance and stiffness after ELAM treatment will allow us to better contextualize this change in diameter.

Here we describe age-related degeneration of the aorta and highlight the critical role of mitochondrial function in driving of aortic aging. Furthermore, our results suggest that a late life treatment paradigm could limit the number of people that require surgery for aortic aneurysms and reduce the burden of diseases like hypertension and heart failure. Thus, targeting the basic mechanisms of aging in the aorta can improve human healthspan and highlights new avenues for investigation.

## Methods

### Experimental Design

This study sought to investigate the role of mitochondria in age-related changes in the aorta. Thus, young (5-6 months) and aged (24-25 months) C57Bl/6JNIA mice either received the mitochondrial-targeted therapeutic elamipretide (ELAM; SS-31) or no treatment for 8 weeks. Mice received pre- and post-treatment blood pressure and echocardiographic measurements to control for inadvertent cardiovascular effects of ELAM. At treatment endpoint, mice were euthanized and thoracic aortas were harvested. The aorta was taken for either tissue respirometry or saved for histologic and RNA analysis.

### Animals

Male and female C57BL/6JNIA mice (C57BL/6J mice maintained at the National Institute on Aging (NIA) aging colony) were acquired from the NIA colony and housed in a specific-pathogen free facility at the University of Washington. All mice were maintained at 21°C on a 14/10 light/dark cycle at 30–70% humidity and given standard mouse chow (LabDiet PicoLab® Rodent Diet 20) and water ad libitum with no deviation prior to or after experimental procedures[67]. This study was reviewed and approved by the University of Washington Institutional Animal Care and Use Committee (IACUC). Similar numbers of males and females were used for all experiments.

### Elamipretide Treatment

Mice were treated with ELAM by osmotic minipumps (Alzet #1004, Cupertino, CA) for 8 weeks at a dose of 3mg/kg/d[12, 67]. Young mice were treated at 5-6 months of age. Aged mice were treated at 24-25 months of age. Prior to surgery, mice underwent baseline echocardiography and non-invasive blood pressure readings. Pumps were implanted in a sterile fashion under isoflurane anesthesia with 0.05mg/kg perioperative subcutaneous buprenorphine analgesia, in accordance with approved IACUC protocol. At four weeks, pumps were exchanged for a freshly loaded pump, implanted in similar fashion as before. Mice randomized to the no treatment arm did not undergo any surgery or treatments. At 8 weeks of treatment, repeat echocardiography and blood pressure readings were performed. ELAM was provided by Stealth BioTherapeutics Inc (Needham, MA). Mice were euthanized for tissue harvest at treatment endpoint.

### Blood Pressure Readings

Non-invasive blood pressure readings were made just prior to pump implantation and treatment endpoint (at 8 weeks of treatment) using the CODA system (Kent Scientific, Torrington, CT). Mice were kept fully awake for the procedure. The machine was used according to the manufacturer’s instructions (kentscientific.com). Fifteen readings were taken per animal, with the first five readings discarded from analysis as the mice acclimated to the holding chamber.

Any further readings with improper waveforms were excluded. The remaining readings were averaged together for analysis.

### Echocardiography

Echocardiography was performed and analyzed on mice just prior to pump implantation and just prior to treatment endpoint (at 8 weeks of treatment) (FUJIFILM VisualSonics, Toronto, Canada). Mice were placed under isoflurane anesthesia with heat support. Parasternal long axis views of the left ventricle were used to assess left ventricular ejection fraction, chamber size, and wall thickness via B-mode. Aortic size was determined with direct longitudinal visualization of the aortic root, ascending aorta, mid-aortic arch, and distal aortic arch, panning the probe to achieve the largest size during systole (increasing the likelihood of measuring the actual diameter). Each measurement was done in triplicate and was remeasured if the standard deviation was >10% of the average.

### Aorta Harvest

The mouse aortas were harvested from root to celiac origin with surrounding vascular structures and fat dissected away. Any aneurysm, dissection, or rupture was noted. For some mice the aorta was taken for tissue respirometry. For other mice the aorta was taken for histology and RNA analysis. Aorta saved for tissue analysis was washed multiple times in 4°C PBS, divided at the distal arch, and a portion of the distal arch was saved for histologic analysis in 10% neutral buffered formalin fixation (NBF). The rest of the aorta was flash frozen in liquid nitrogen and stored at −80°C.

### Tissue Respirometry

Tissue respirometry measurements were made with an Oxygraph 2K respirometer (Oroboros Instruments, Innsbruck, Austria) at 37°C in 0.5 mL chambers. Aorta taken for tissue respirometry was divided at the distal arch into a proximal ascending/arch portion and a distal descending thoracic aorta (DTA) portion. The aorta was then washed in 4°C isolation buffer (IB: 20 mM CaK_2_EGTA, 20 mM K_2_EGTA, 6.56 mM MgCl_2_, 0.5 mM DTT, 50 mM K-MES, 20 mM imidazole, 20 mM taurine, 5.77 mM Na_2_ATP, and 15 mM phosphocreatine) and carefully evaluated under a dissection microscope to remove any residual surrounding fat. The aorta was then divided at the distal arch, blotted until dry, and weighed. Following this, the aorta was permeabilized for 40 minutes in a solution of IB and 50 μg/ml of saponin. Afterwards, the aorta was washed twice in respiration buffer (RB: 0.5 mM EGTA, 3 mM MgCl_2_-6H_2_O, 10 mM KH_2_PO_4_, 20 mM HEPES, 110 mM sucrose, 100 mM mannitol, 60 mM K-MES, 20 mM taurine, 1 g/L BSA, pH 7.1) for ten minutes each. The ascending/arch and DTA were placed in separate 0.5 mL respirometer chambers, each hyper-oxygenated to 420-500 μM O_2_. Reagents were prepared and administered according to Oroboros instructions (bioblast.at). Reagents were added in the following order: simultaneous 10 mM glutamate and 0.5 mM malate (leak state), 150 μM ADP (sub-saturating ADP concentration, state III respiration), 2.5 mM ADP (saturating ADP concentration), 5 mM pyruvate (complex I (CI)), 10 mM succinate (maximal coupled respiration), 0.5 μM carbonyl cyanide-p-trifluoromethoxyphenylhydrazone (FCCP) (maximal uncoupled respiration), 0.5 μM rotenone (complex II (CII)), 2.5 μM antimycin A (background consumption). Oxygen consumption rate (OCR, pmol/sec/mg tissue) was recorded with each substrate and subsequently normalized to the OCR after antimycin A administration. At assay completion, the aortic tissue was flash frozen in liquid nitrogen and stored at −80°C.

### Histologic Assessment

Aortic tissue fixed in NBF was processed, sectioned, and stained with either hematoxylin and Van Gieson’s stain (for elastin visualization) or Masson’s Trichrome stain (for collagen visualization) by the University of Washington Histology & Imaging Core. Three sections for each tissue sample were analyzed under a high-powered field at 10x and subsequently averaged together. Image analysis was done using ImageJ (National Institutes of Health, Bethesda, MD). Media thickness was determined as the distance from the internal elastic lamina to the external elastic lamina, using 12 measurements around the aorta in a clockwise fashion[78]. The number of elastin breaks were also counted for each section, normalized to the internal elastic laminar circumference[78]. Proportion of collagen content was determined by outlining the aortic media, defining the stained collagen color thresholds (Hue 115-195, Saturation 25-255, Brightness 20-255), converting the image into greyscale, and measuring the proportion of stained area[79].

### RNA isolation, RT-qPCR, and RNA sequencing

RNA isolation was done using the QIAgen RNEasy kit (#74104 QIAGEN, Hilden, Germany). Aorta kept at −80 °C was thawed to 4°C, residual surrounding fat was dissected at 4°C, and the aorta was pestle homogenized in RLT buffer (#74104 QIAGEN, Hilden, Germany) supplemented with 40 mM DTT. The rest of the isolation was per the kit protocol. RNA concentration and quality was determined using a NanoDrop One spectrophotometer (ThermoScientific, Waltham, MA). RT-qPCR was performed using the Verso 1-Step Reaction Mix (#AB4101A, ThermoScientific) with a final reaction volume of 15 μL. Primers used were: p16/CDKN2a (Thermo Fisher #4331182, assay ID Mm01257348_m1), p21/CDKN1a (Thermo Fisher #4331182, assay ID Mm04205640_g1), MMP9 (Thermo Fisher #4331182, assay ID Mm00442991_m1), and GAPDH (Thermo Fisher #4331182, assay ID Mm99999915_g1).

GAPDH was used as a housekeeping gene. Aliquots of at least 450ng of RNA were sent to Azenta Life Sciences (Burlington, MA) for bulk RNA sequencing.

### Transcriptome Analysis

The RT-qPCR data were analyzed in a standard ddCt method. Ct values for genes in each sample were normalized to GAPDH Ct, and then ddCt values were obtained by finding the difference between the dCt of each sample relative to the average of the dCt values in the Young NT group. Statistical analysis was performed on ddCt values, and plots were obtained using relative expression values (2^−ddCt^).

The RNA-seq data were the log-normalized expression of 21,326 mouse genes among 24 samples that had been adjusted for technical variation by Azenta. After filtering for missing values, we analyzed the remaining 20,607 genes measured in all samples. We performed PCA on the data after scaling the expression of each gene and used the Tracy-Widom test to determine the significance of each PC[80]. For univariate analysis, we modeled the expression of each gene as a function of treatment (NT and ELAM), age as two groups (old and young), and the interaction between treatment and age, along with fixed effect of sex. We used ANOVA and false-discovery rate (FDR) adjustment to test the effect (*F*>0) of sex, treatment, age, and treatments x age interaction.

The SenMayo gene set was retrieved from www.gsea-msigdb.org, and 114 of the 125 SenMayo annotations were matched to the *Mus musculus ENSEMBL Gene IDs* in the RNA-seq data. Enrichment of SenMayo genes was tested by Fisher’s Exact test. Enrichment of GO terms among the genes was assessed using the clusterProfiler package, using FDR to adjust for multiple testing, and the rrvgpo package to reduce GO terms by redundancy[81, 82].

### Statistical Analysis

All data presented are mean +/- standard deviation (except box-and-whisker plots, which present the mean and range). Data were visualized prior to analysis to evaluate for obvious skewing/non-normality. Comparisons between two groups were performed either with student’s t-test or multiple t-tests with a Holm-Sidak correction. Comparisons between all four treatment groups (young NT, young ELAM, aged NT, and aged ELAM) were done with a 2-way ANOVA followed by Fisher’s least significant difference for multiple comparisons (given the inclusion of the age-treatment interaction term). Transcriptomic analysis is described above.

## Author Contributions

- Conceptualization: AD, DJM
- Methodology: AD, GP, BH, AMH, RS, DJM
- Investigation: AD, BH, AMH, RS
- Visualization: AD
- Supervision: DJM
- Writing—original draft: AD
- Writing—review & editing: AD, GP, BH, ST, SD, CRB, BH, JDP, MSM, DJM

## Data and Materials Availability

- All data needed to evaluate the conclusions in the paper are available in the main text or the supplementary materials

## Supplemental Figures

**Supplemental Figure 1:**
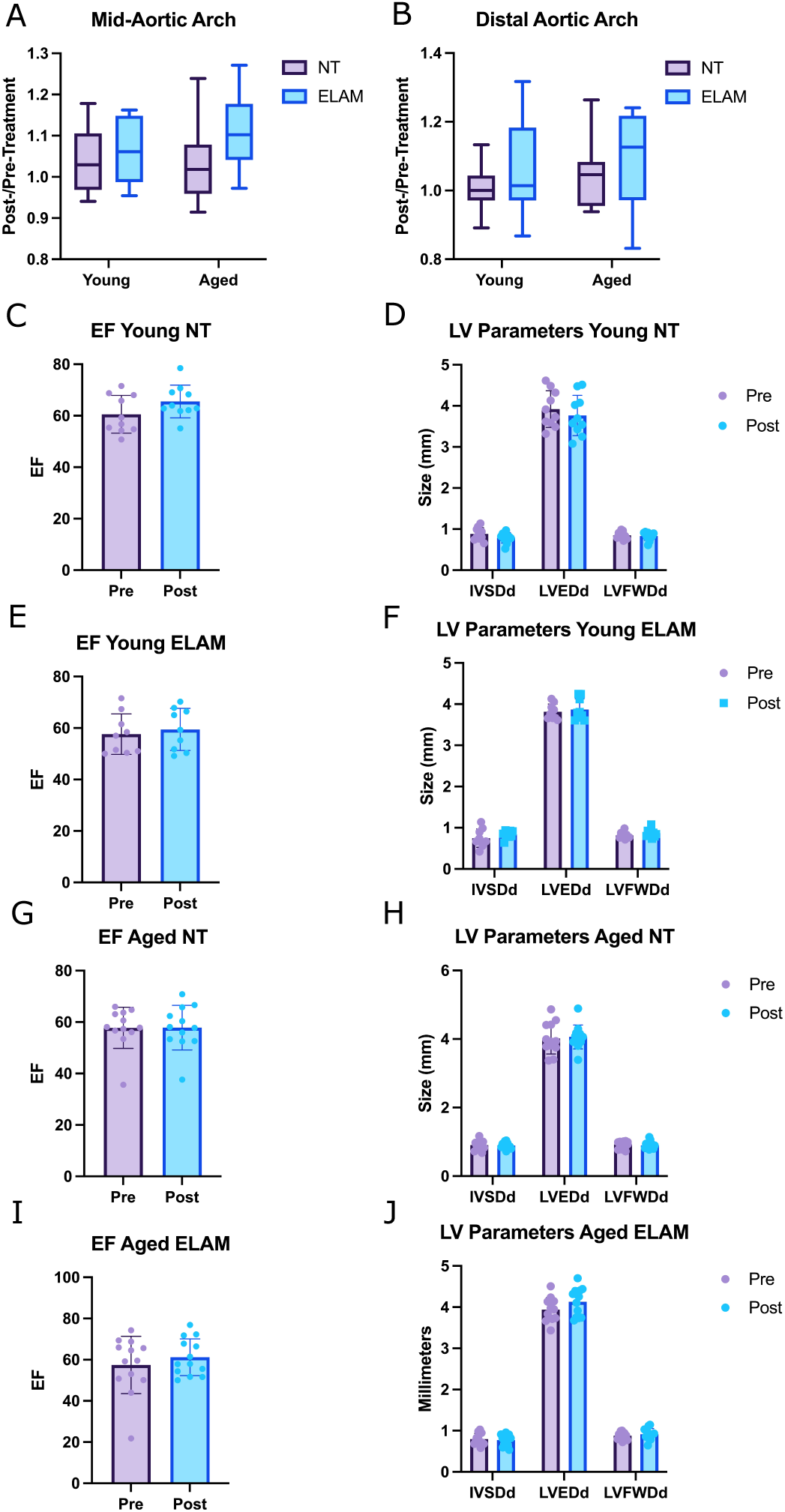
Echo parameters for each treatment group. Relative change of the mid-arch (A) and distal arch (B) sizes over the 8 week treatment period. Ejection fraction (EF) and left ventricular (LV) dimensions for the young NT (C-D), young ELAM (E-F), aged NT (G-H), and aged ELAM (I-J) groups. IVSDd—interventricular septal thickness during diastole; LVEDd—left ventricular end-diastolic diameter; LVFWDd—left ventricular free wall thickness during diastole. (A-B) 2-way ANOVA, Fisher’s least significant difference. (C,E,G,I) Student’s t-test. (D,F,H,J) Multiple t-tests, Holm-Sidak correction. * p<0.05, ** p<0.01, *** p<0.001, **** p<0.0001.

**Supplemental Figure 2:**
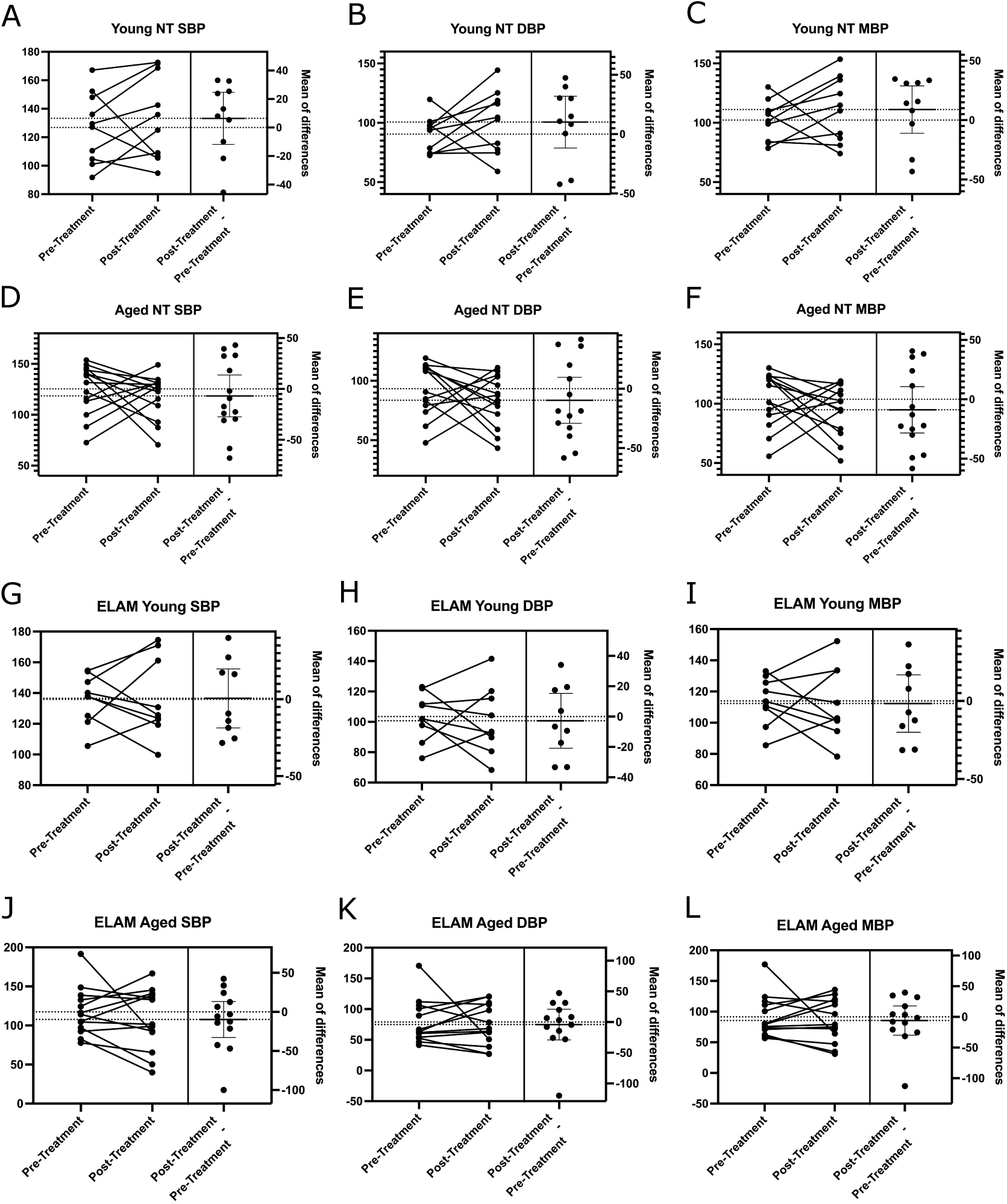
Blood pressure change over the study period. Pre-treatment and post-treatment systolic blood pressure (SBP), diastolic blood pressure (DBP), and mean blood pressure (MBP) were obtained for the young NT (A-C), aged NT (D-F), young ELAM (G-I), and aged ELAM (J-L) groups. Each graph shows the paired changes in blood pressure over the study period for the treatment groups. In the line plot, pre-treatment BP is on the left, post-treatment BP is on the right. The right-most plot in each graph shows the difference over the treatment period for each mouse (post-treatment BP – pre-treatment BP). Both the mean and zero are shown with dotted horizontal lines. ELAM had no effect on BP. Paired student’s t-test, * p<0.05, ** p<0.01, *** p<0.001, **** p<0.0001.

**Supplemental Figure 3:**
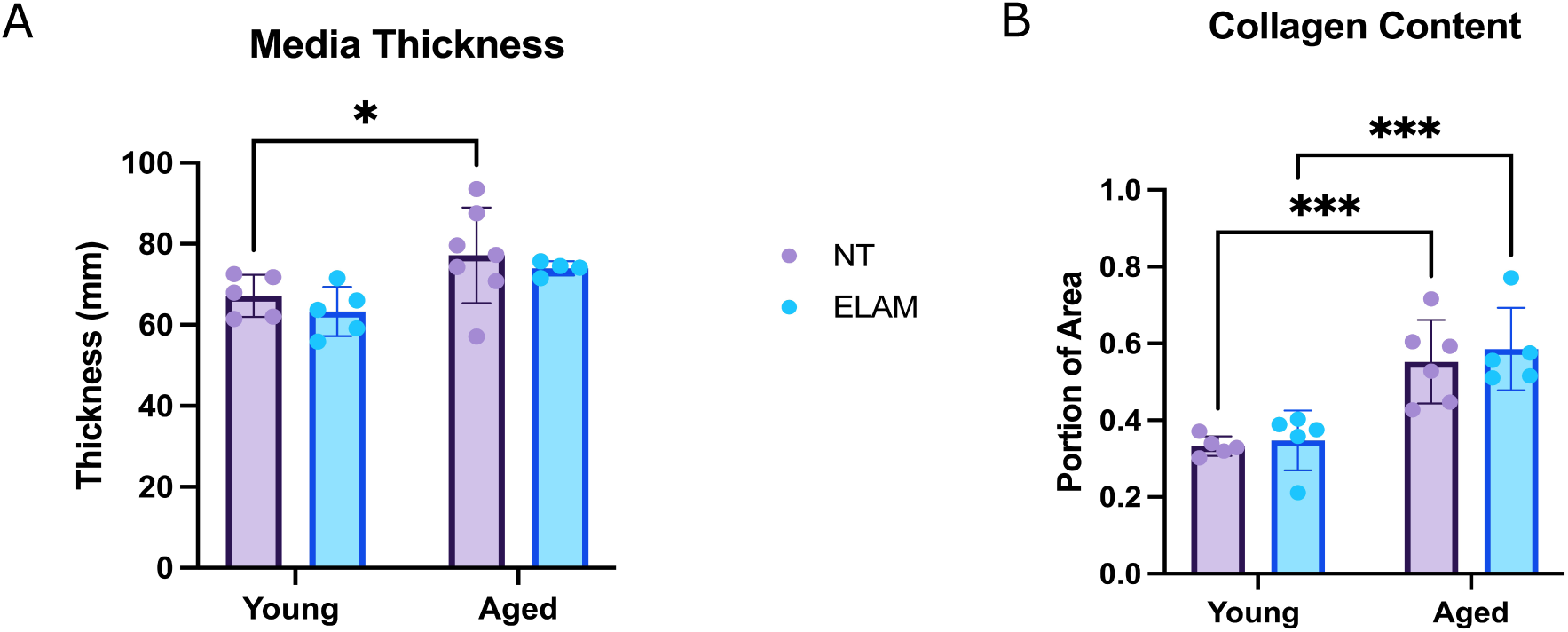
ELAM treatment did not alter media thickness or collagen content. Histologic assessment of media thickness (A) and collagen content (B) in the four treatments groups (n=5-6 per group) revealed no difference with ELAM treatment. 2-way ANOVA, Fisher’s least significant difference, * p<0.05, ** p<0.01, *** p<0.001, **** p<0.0001.

**Supplemental Figure 4:**
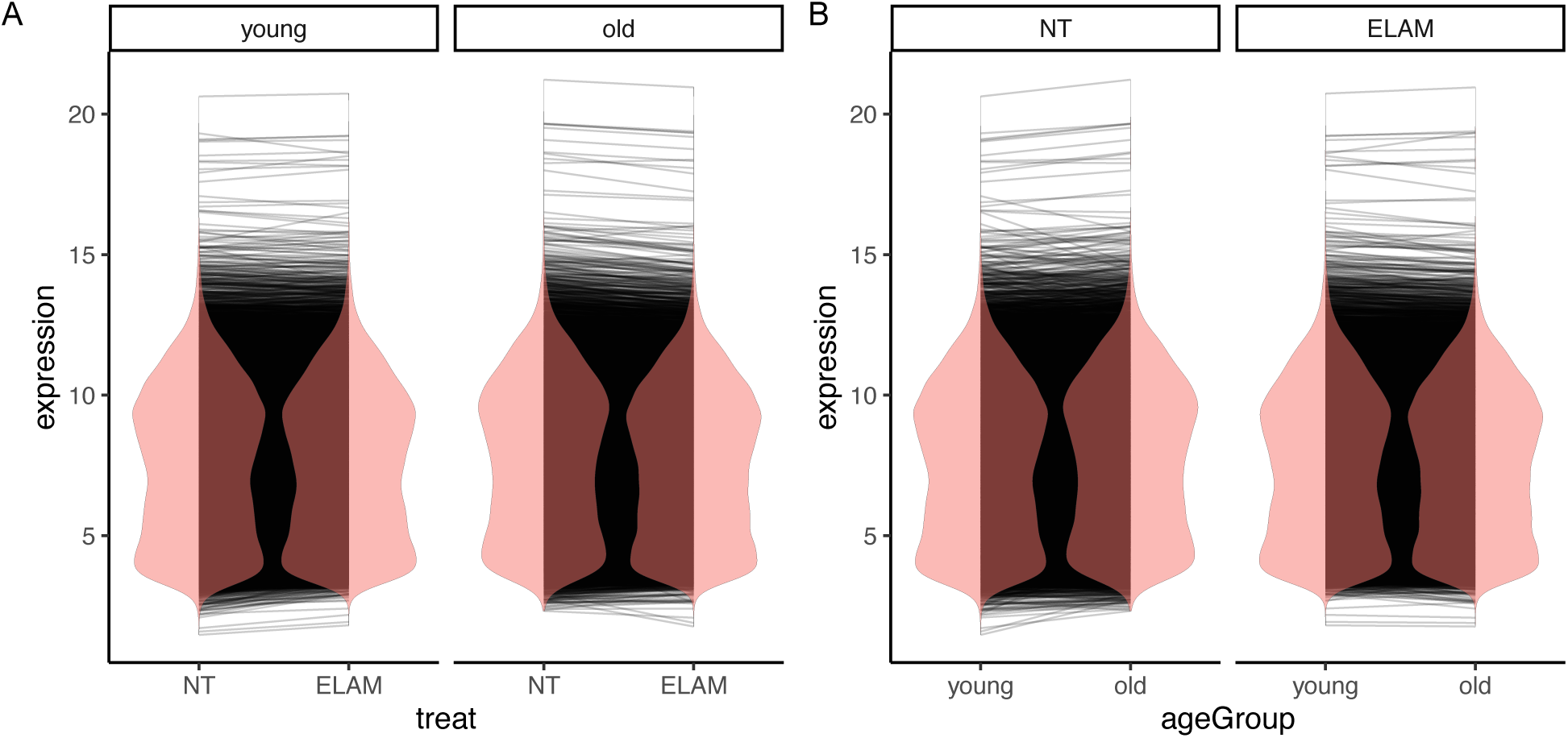
Differences in expression levels in genes with a significant age by treatment interaction. The mean background expression level of genes with a significant age by treatment interaction in NT and ELAM treated mice are connected by lines in (A). The mean background expression level of genes with a significant age by treatment interaction in young and aged mice are connected by lines in (B). ANOVA with false-discovery rate (FDR) adjustment (*F*>0).

## Supplemental Tables

**Supplemental Table 1:** All terms assessed for effects on gene expression. ANOVA and false-discovery rate (FDR) adjustment (<5%).

| Term | # genes affected (FDR <5%) |
| --- | --- |
| treatment | 6,287 |
| age (young vs. old) | 6,061 |
| treatment x age | 4,339 |
| sex | 12 |

## Supplemental Data

Please see Supplemental Data excel doc (SuppDat1)

**Supplemental Data 1: GO terms significantly enriched by genes with age-treatment interaction**. Terms were broadly organized into mitochondrial (1), metabolic (2), inflammatory (3), redox (4), and other cellular function categories (5).

